# Calcium dynamics of skin-resident macrophages during homeostasis and tissue injury

**DOI:** 10.1101/2024.09.24.614510

**Authors:** Pearl A. Leon Guerrero, Jeffrey P. Rasmussen, Eric Peterman

## Abstract

Immune cells depend on rapid changes in intracellular calcium activity to modulate cell function. Skin contains diverse immune cell types and is critically dependent on calcium signaling for homeostasis and repair, yet the dynamics and functions of calcium in skin immune cells remain poorly understood. Here, we characterize calcium activity in Langerhans cells, skin-resident macrophages responsible for surveillance and clearance of cellular debris after tissue damage. Langerhans cells reside in the epidermis and extend dynamic dendrites in close proximity to adjacent keratinocytes and somatosensory peripheral axons. We find that homeostatic Langerhans cells exhibit spontaneous and transient changes in calcium activity, with calcium flux occurring primarily in the cell body and rarely in the dendrites. Triggering somatosensory axon degeneration increases the frequency of calcium activity in Langerhans cell dendrites. By contrast, we show that Langerhans cells exhibit a sustained increase in intracellular calcium following engulfment of damaged keratinocytes. Altering intracellular calcium activity leads to a decrease in engulfment efficiency of keratinocyte debris. Our findings demonstrate that Langerhans cells exhibit context-specific changes in calcium activity and highlight the utility of skin as an accessible model for imaging calcium dynamics in tissue-resident macrophages.

**SIGNIFICANCE STATEMENT:**

- Calcium activity in immune cells is thought to regulate cell function, but studies focusing on tissue-resident macrophages are limited.
- Skin-resident macrophages known as Langerhans cells exhibit rapid and transient changes in calcium activity in homeostatic conditions, which can change depending on the type of tissue injury inflicted. Pharmacological perturbation of calcium activity leads to a decrease in Langerhans cell engulfment.
- These findings suggest calcium activity is important for tissue surveillance by Langerhans cells.

## INTRODUCTION

Diverse immune cell types patrol our bodies to eliminate foreign pathogens, clear cellular debris and maintain tissue homeostasis. Immune cell function requires dynamic and rapid responses to environmental signals encountered by transmembrane receptors. Among the signaling cascades elicited upon receptor engagement within immune cells, calcium serves as an important secondary messenger in a variety of contexts such as antigen recognition and wound healing (Ghilardi et al., 2020; Trebak and Kinet, 2019).

As a superficial neuroimmune organ that undergoes constant turnover during homeostasis and rapid remodeling after injury, skin is an excellent model to study calcium function and dynamics. Calcium signaling promotes normal keratinocyte function, including differentiation (Elias et al., 2002; Hennings et al., 1980; Yuspa et al., 1989), proliferation (Moore et al., 2023), and epithelial barrier maintenance (Celli et al., 2021). Calcium is also required for cutaneous regeneration in multiple *in vivo* models (Balaji et al., 2017; Han et al., 2024; Moore et al., 2023; Restrepo and Basler, 2016; Varadarajan et al., 2022; Xu and Chisholm, 2011). Altered calcium signaling is implicated in several cutaneous disorders, including atopic dermatitis and psoriasis (Wang et al., 2021). Despite the importance of calcium in skin homeostasis, repair, and pathologies, little is known about the dynamics and functional roles of calcium in skin-resident immune cells. This is in part due to a lack of immune cell diversity in invertebrate systems and technical challenges of *in vivo* imaging in mammalian systems.

Zebrafish contains a diverse repertoire of immune cells and has optically clear skin, making it an excellent system to study cutaneous immune cell behaviors. Similar to human skin, the adult zebrafish epidermis contains stratified layers of keratinocytes that interact with the peripheral endings of somatosensory neurons and various immune cell types (Ferrero et al., 2020; Hui et al., 2017; Lin et al., 2019; Lugo-Villarino et al., 2010; Rasmussen et al., 2018; Sire et al., 1997; Wittamer et al., 2011; Zhou et al., 2023). To gain better insight into immune cell dynamics in skin, we recently developed models of axon and keratinocyte damage in adult zebrafish skin (Peterman et al., 2023; Peterman et al., 2024). Using these damage models, we found that Langerhans cells, an epidermal-resident immune cell type in vertebrate skin (Doebel et al., 2017), exhibit macrophage-like behaviors, including the engulfment of degenerating axonal debris and laser-damaged keratinocytes (Peterman et al., 2023; Peterman et al., 2024). Previous studies concluded that in vitro exposure to capsaicin can raise Langerhans cell calcium levels (Mariotton et al., 2023) and treatment with calcium channel inhibitors affects the morphology and antigen presentation abilities of Langerhans cells (Diezel et al., 1989; Katoh et al., 1997). However, whether Langerhans cells dynamically alter calcium levels in their native skin environment remains unknown. Here, we use our accessible experimental platform to describe and perturb the calcium dynamics of Langerhans cells.

## RESULTS AND DISCUSSION

### Langerhans cells exhibit sporadic, unsynchronized calcium dynamics in homeostatic conditions

Imaging immune cell calcium dynamics in living tissue is technically challenging but has precedent in other tissue-resident macrophages (Hughes and Appel, 2020; Mehari et al., 2022; Umpierre et al., 2020). Zebrafish scales are bony dermal appendages arranged in a shingle-like pattern on the zebrafish trunk, with the epidermis and resident cells (such as Langerhans cells) overlaid on the scale surface (Aman and Parichy, 2024). The superficial location and flat structure of scales make them amenable to live imaging (Cox et al., 2018; De Simone et al., 2021). To observe calcium dynamics in Langerhans cells, we used the *mpeg1*.*1* promoter to label skin macrophages, including Langerhans cells, in adult zebrafish (He et al., 2018; Kuil et al., 2020; Lin et al., 2019; Peterman et al., 2023; Peterman et al., 2024). We generated double-transgenic fish expressing *Tg(mpeg1*.*1:GCaMP6s-CAAX)* (Hughes and Appel, 2020) and *Tg(mpeg1*.*1:mCherry)* (Ellett et al., 2011), and used the membrane-localized calcium biosensor GCaMP6s-CAAX to follow calcium dynamics and cytosolic mCherry to visualize the entirety of the Langerhans cell body. To determine whether Langerhans cells exhibited calcium activity during homeostasis, we performed *in vivo* confocal microscopy on anesthetized adult fish to image trunk skin every 5 seconds for 15 minutes **(Figure 1A, B)**. We observed transient elevations in calcium levels, suggesting that Langerhans cells exhibit spontaneous calcium dynamics **(Figure 1C)**.

**Figure 1.**
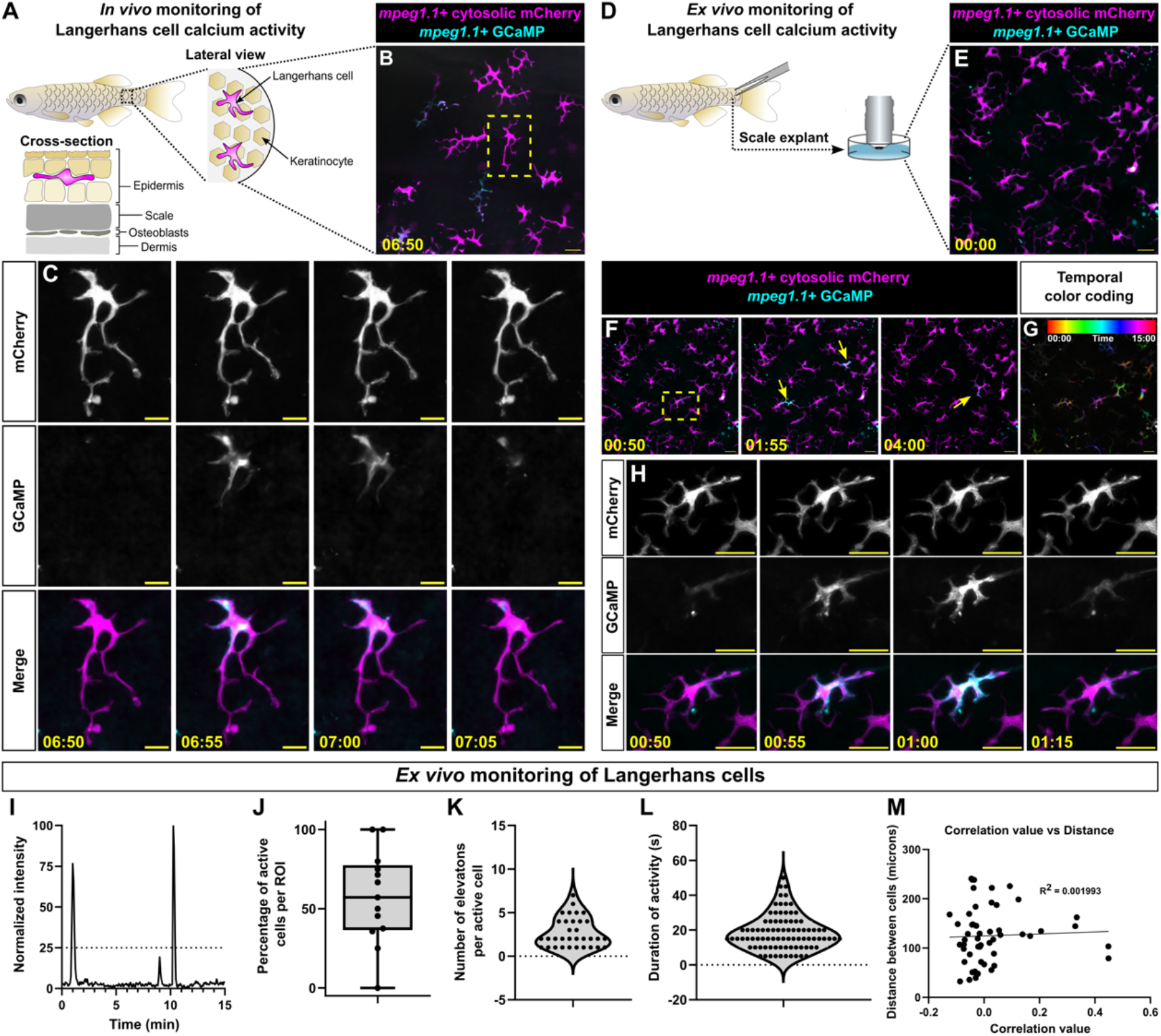
Langerhans cells elevate calcium levels frequently and transiently. **A**. Schematic of *in vivo* imaging paradigm. **B**. Confocal micrograph from *in vivo* time lapse imaging showing *Tg(mpeg1*.*1:mCherry;mpeg1*.*1:GCaMP6s-CAAX)*+ Langerhans cells. Yellow box denotes the ROI magnified in **(C).C**. Inset still images showing Tg(*mpeg1*.*1*:mCherry;*mpeg1*.*1*:GCaMP6s-CAAX)+ Langerhans cell exhibiting increase in calcium activity. **D**. Schematic of *ex vivo* imaging paradigm. **E**. Confocal micrograph from *in vivo* time lapse imaging showing *Tg(mpeg1*.*1:mCherry;mpeg1*.*1:GCaMP6s-CAAX)*+ Langerhans cells. **F**. Confocal micrographs from *in vivo* time lapse imaging showing *Tg(mpeg1*.*1:mCherry;mpeg1*.*1:GCaMP6s-CAAX)*+ Langerhans cells exhibiting changes in calcium levels. Yellow arrows indicate cells exhibiting increases in calcium levels. Yellow box denotes the ROI magnified in **(H). G**. Temporal color coding of the 15 minute time-lapse session from **F**, indicating that multiple Langerhans cells exhibited increases in calcium activity. **H**. Still images showing a *Tg(mpeg1*.*1:mCherry;mpeg1*.*1:GCaMP6s-CAAX)*+ Langerhans cell exhibiting an increase in calcium activity. **I**. Normalized intensity plot of GCaMP signal in Langerhans cell shown in **(H)**. A minimum threshold of 25% normalized intensity (dashed line) was used to score cells as positive for activity (see Materials and Methods). **J**. Boxplot of the percentage of cells exhibiting calcium flux during the 15 minute imaging window, *n* = 13 ROIs tracked from *N* = 3 fish. **K**. Violin plot showing the number of calcium elevations an individual cell produced within the 15 minute imaging window, *n* = 33 cells tracked from *N* = 13 ROIs. **L**. Violin plot showing the duration of elevated calcium levels, *n* = 87 active cells tracked from *N* = 13 ROIs. **M**. Scatter plot of correlation of time when cells exhibited increases in calcium levels versus distance from other Langerhans cells, *n* = 52 correlation plots tracked from *N* = 9 ROIs. Time stamps denote mm:ss. Scale bars: 20 μm (B, E, F, H), 10 μm (C).

To facilitate image acquisition and experimental manipulation, we turned to our previously established skin explant system (Peterman et al., 2023; Peterman et al., 2024). We used confocal microscopy to image scale explants **(Figure 1D, E)** and, as we observed *in vivo*, we found that Langerhans cells displayed transient fluctuations in calcium levels **(Figure 1F-I, Supplemental Movie 1**, see *Materials and Methods* for quantification details). The majority of cells (57.2%) showed changes in calcium levels in a given ROI over our 15 minute imaging window **(Figure 1J)**. Of the cells that showed changes in activity, 69.7% displayed multiple events, with an average of 2.7 calcium elevation events in the 15 minute window **(Figure 1K)**. The average length of elevated calcium levels was 18.56 s **(Figure 1L)**. In certain cell types, such as basal keratinocytes, increases in calcium levels can propagate between cells across a tissue (Moore et al., 2023). To infer if the changes in calcium levels of Langerhans cells propagated across the epidermis, we compared when cells exhibited increases in calcium levels with their distance from each other. However, we did not find a significant correlation, suggesting the changes in intracellular calcium levels are unrelated to the calcium activity of neighboring Langerhans cells **(Figure 1M)**.

Localized calcium activity in microglial dendrites precedes engulfment of myelin debris in zebrafish embryos (Hughes and Appel, 2020). When we quantified where calcium activity occurred in homeostatic Langerhans cells, we noted that elevated calcium levels most often occurred throughout the entire cell (89.9%; **Figure 2A, C, Supplemental Video 2)**, rather than within individual dendrites (10.1%; **Figure 2B, C, Supplemental Video 2)**. Additionally, whole cell calcium activity lasted significantly longer than dendrite only activity **(Figure 2D)**. Altogether, our results suggest Langerhans cells exhibit sporadic, unsynchronized changes in intracellular calcium levels in homeostatic conditions, with the majority of the calcium elevation events occurring throughout the cell.

**Figure 2.**
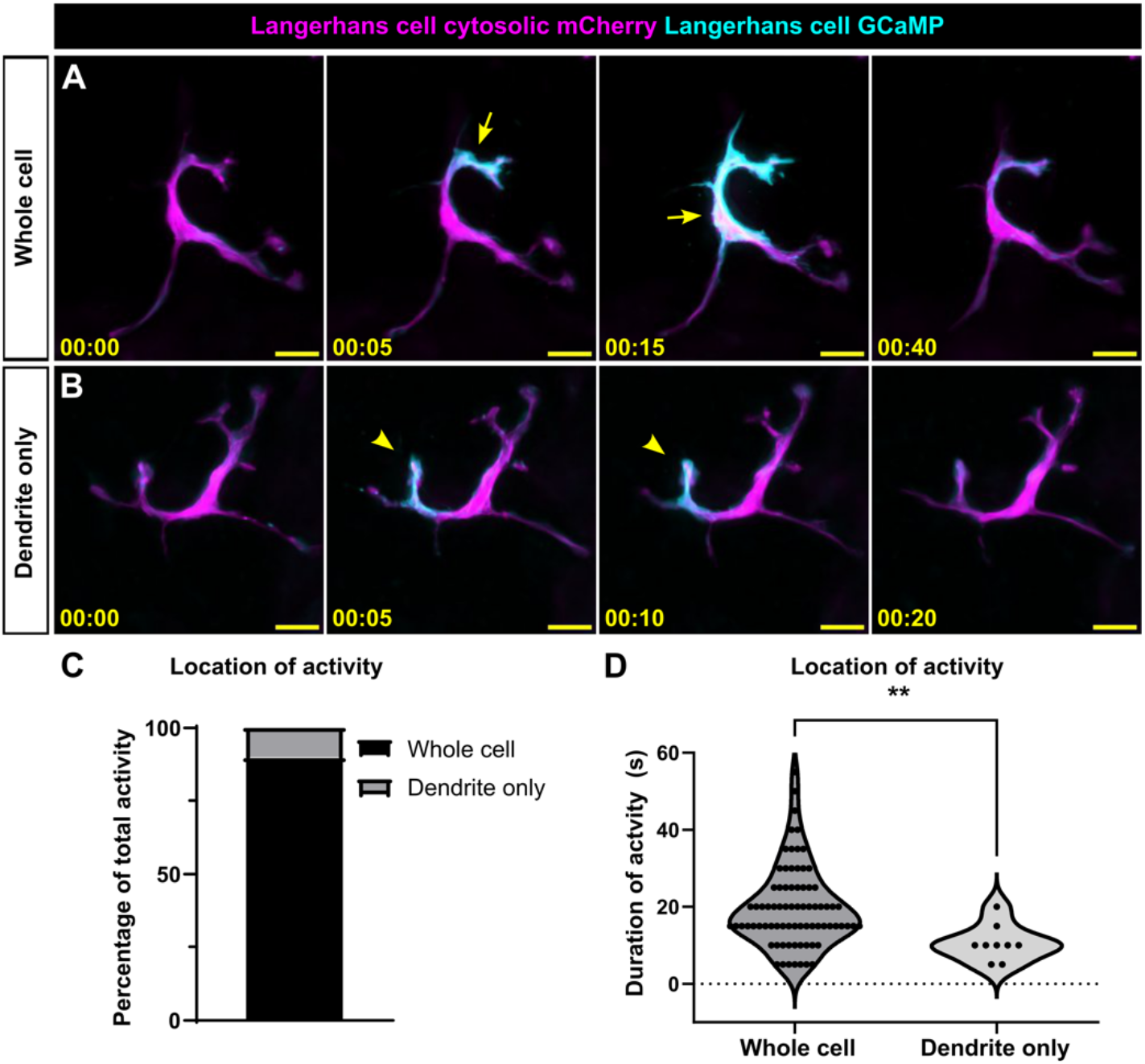
Spatial heterogeneity of calcium activity in Langerhans cells. **A, B**. Confocal micrographs from time lapse imaging showing a *Tg(mpeg1*.*1:mCherry;mpeg1*.*1:GCaMP6s-CAAX)*+ Langerhans cell exhibiting calcium activity in the whole cell **(A)**, or the dendrite only **(B)**. Arrows in **A** point to whole cell activity, arrowheads in **B** point to dendrite only activity. **C**. Quantification of the percentage of calcium activity occurring in the whole cell or dendrite only, *n* = 88 cells tracked from 13 ROIs from *N* = 3 fish. **D**. Violin plots showing the duration of elevated calcium in the whole cells and dendrites, *n* = 87 calcium elevation events tracked from *N* = 34 cells. Mann-Whitney U test was used to determine significance in **(D)**, ** = *p* < 0.01. Time stamps denote mm:ss. Scale bars: 10 μm (A), 20 μm (B).

### Axon degeneration alters the subcellular location of calcium dynamics

Phagocytosis and phagosome formation are among the immune cell functions regulated by changes in calcium levels (Westman et al., 2019). Thus, we next sought to understand if tissue damage elicited changes in Langerhans cell calcium activity. To this end, we first tracked Langerhans cell calcium dynamics during axon degeneration. Somatosensory axons emanating from dorsal root ganglia innervate the scale epidermis (Rasmussen et al., 2018). Removing scales triggers axon degeneration; after 4-6 hours, axons undergo Wallerian degeneration and produce axonal debris (1-3 μm in diameter), which Langerhans cells engulf (Peterman et al., 2023). When compared to calcium activity in homeostatic conditions, Langerhans cells exhibited similar calcium dynamics upon axon degeneration and subsequent debris engulfment **(Figure 3A, B, D)**. Intriguingly, we observed an increase in the percentage of dendrite-only events, up to 34.9 percent (from 10.1 percent) (**Figure 3C)**. These data suggest that engulfment of small axonal debris does not appreciably affect calcium dynamics but does change the spatial location of calcium activity.

**Figure 3.**
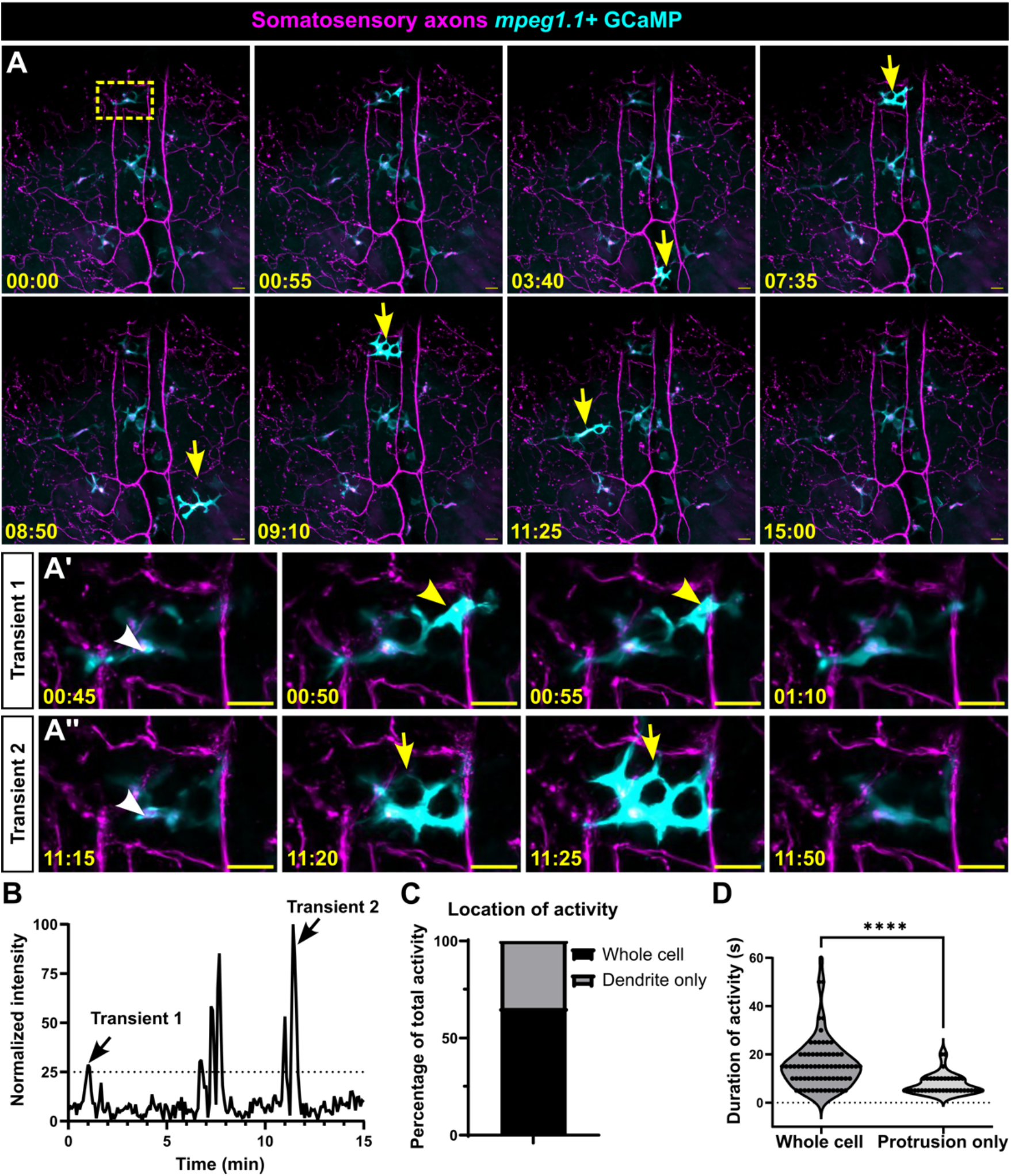
Location, not duration, of calcium activity in Langerhans cells is altered during axon degeneration. **A**. Confocal micrographs from time lapse imaging showing *Tg(p2rx3a:mCherry)*+ axons degenerating and *Tg(mpeg1*.*1:GCaMP6s-CAAX)*+ Langerhans cells engulfing debris and exhibiting changes in calcium levels. Yellow box in first frame denotes insets in **A’, A’’**. White arrowheads in **A’, A’’** indicate engulfed axonal debris. Yellow arrows indicate whole cell activity, yellow arrowheads indicate dendrite-only activity. **B**. Intensity plot of GCaMP signal in Langerhans cell from panels **A**’, **A’’. C**. Quantification of the percentage of calcium activity occurring in the whole cell or dendrite only, *n* = 109 calcium elevation events tracked from *N* = 26 cells. **D**. Violin plots showing the duration of elevated calcium in the whole cells and dendrites, *n* = 109 calcium elevation events tracked from *N* = 26 cells. Mann-Whitney U test was used to determine significance in **(D)**, **** = *p* < 0.0001. Timestamps denote mm:ss. Scale bars: 20 μm (A, A’, A’’).

### Keratinocyte debris engulfment triggers prolonged elevation of intracellular calcium levels

Previously, we demonstrated that Langerhans cells undergo a ramified-to-rounded shape transition to accommodate engulfment of keratinocyte debris (10-20 μm in diameter) (Peterman et al., 2024). We next questioned if this shape transition and engulfment method displayed different calcium activity when compared to engulfment of axon debris. We used a pulsed laser to ablate individual keratinocytes, which triggered subsequent engulfment by Langerhans cells. Upon keratinocyte engulfment, 100% of Langerhans cells exhibited high and sustained calcium levels **(Figure 4A, B**, blue line, *n*=12/12, **Supplemental Video 3)**. Notably, Langerhans cells sustained high calcium levels for an average of 61.33 minutes, substantially longer than those observed during homeostasis or axon debris engulfment **(Figure 4C)**. Nearby cells that did not engulf debris did not exhibit sustained elevations of calcium **(Figure 4B**, pink line), suggesting that the increases in calcium levels were due to debris engulfment rather than a non-specific effect of the pulsed laser. In contrast to the sharp drop in calcium activity observed in homeostatic conditions, calcium signal gradually decreased to baseline post-engulfment **(Figure 4B)**. To determine when in the phagocytic process calcium activity increased, we measured the time between phagocytic cup closure and increased calcium levels. We found a heterogenous timing of calcium activity onset; 7/12 cells increased calcium levels within 12 minutes post-engulfment, whereas 5/12 cells increased calcium levels with a delayed onset **(Figure 4D)**. These data suggest that keratinocyte engulfment elicits a long-lasting rise in intracellular calcium within Langherhans cells, although the onset and duration of the elevated calcium varies.

**Figure 4.**
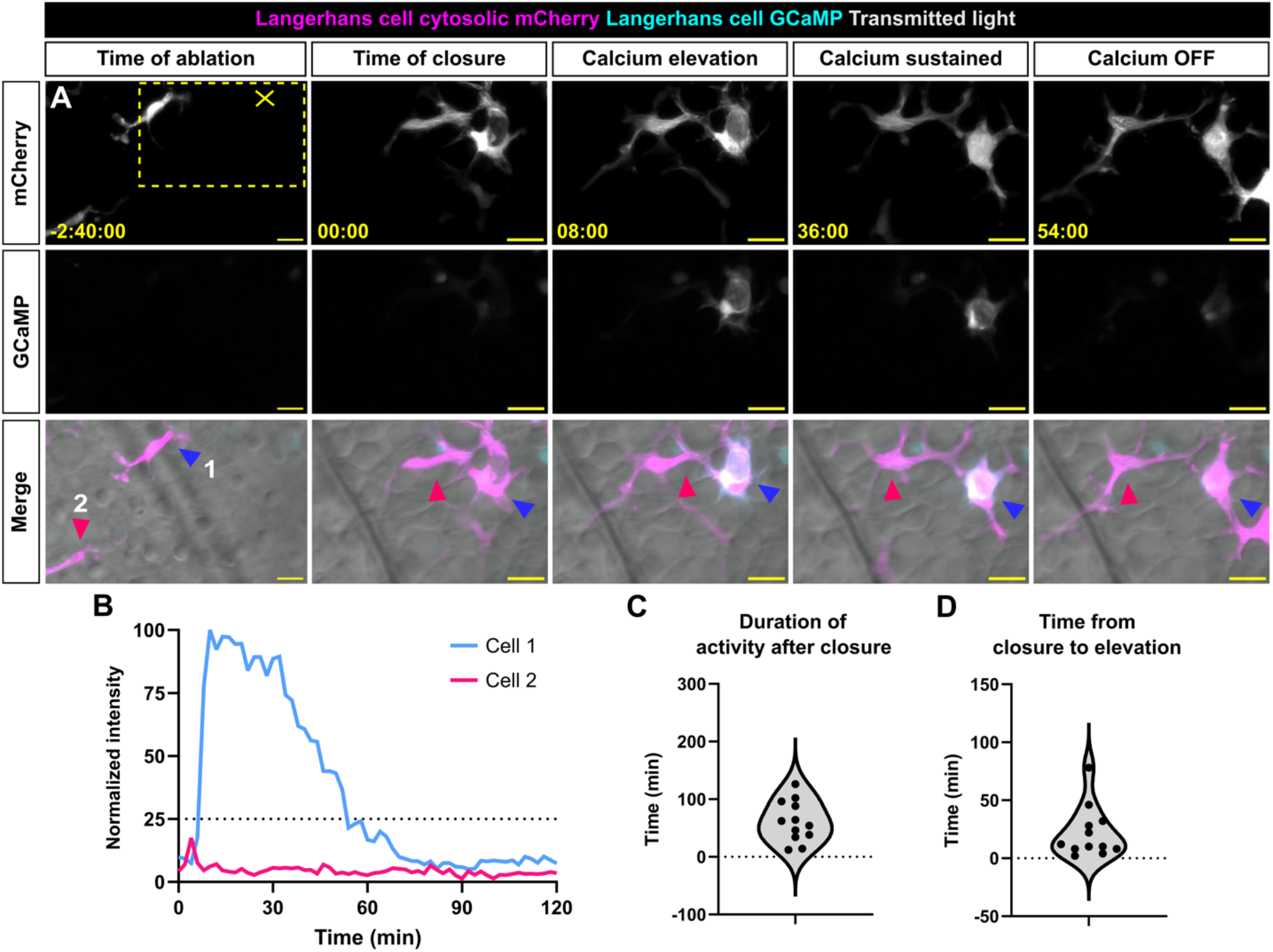
Langerhans cells exhibit elevated and sustained calcium levels after engulfment of keratinocyte debris. **A**. Confocal micrographs from time lapse imaging showing *Tg(mpeg1*.*1:mCherry;mpeg1*.*1:GCaMP6s-CAAX)*+ Langerhans cell engulfing debris after laser ablation of nearby keratinocyte. Yellow dotted box indicates insets used in panels (00:00 to 54:00). Yellow X indicates the site of laser ablation. Blue and pink arrowheads indicate cells tracked in **(B). B**. Intensity plot of GCaMP signal in Langerhans cell from panel **(A). C**. Violin plot showing the length of time from phagocytic cup closure to 25% GCaMP signal, *n* = 12 cells tracked from *N* = 7 scales. **D**. Violin plot showing the duration of >25% GCaMP signal, *n* = 12 cells tracked from *N* = 7 scales. Timestamps denote mm:ss. Scale bars: 20 μm (A).

### Elevated cytosolic calcium levels inhibit debris engulfment

Since Langerhans cells displayed sustained calcium activity during engulfment of keratinocyte debris, we questioned if altering cytosolic calcium levels could impact the efficiency of debris engulfment. Treating skin explants with the calcium chelators BAPTA-AM or EGTA or the ionophore ionomycin led to inconsistent or off-target effects, including widespread cell death. We also observed that treatment with DMSO alone led to increases in Langerhans cell calcium activity. Thus, we turned to thapsigargin, a sarcoplasmic/endoplasmic reticulum calcium ATPase (SERCA) inhibitor soluble in ethanol. Cells typically maintain low cytosolic calcium levels due to electrochemical gradients and the energy-dependent action of SERCA that promotes calcium re-entry into the endoplasmic reticulum (ER) (Westman et al., 2019). Following depletion of ER calcium stores during an event such as phagocytosis, calcium re-enters the ER via SERCA channels. By preventing calcium re-entry to the ER via SERCA inhibition, ER stores become depleted and cytosolic calcium levels remain elevated. We first confirmed that thapsigargin treatment caused elevated calcium levels. Whereas ethanol-treated Langerhans cells displayed normal, transient calcium flashes **(Figure 5A, C, D)**, thapsigargin-treated Langerhans cells exhibited sustained levels of high calcium, often starting 10 minutes post-treatment and persisting for the rest of the 30 minute time lapse **(Figure 5B-D)**.

**Figure 5.**
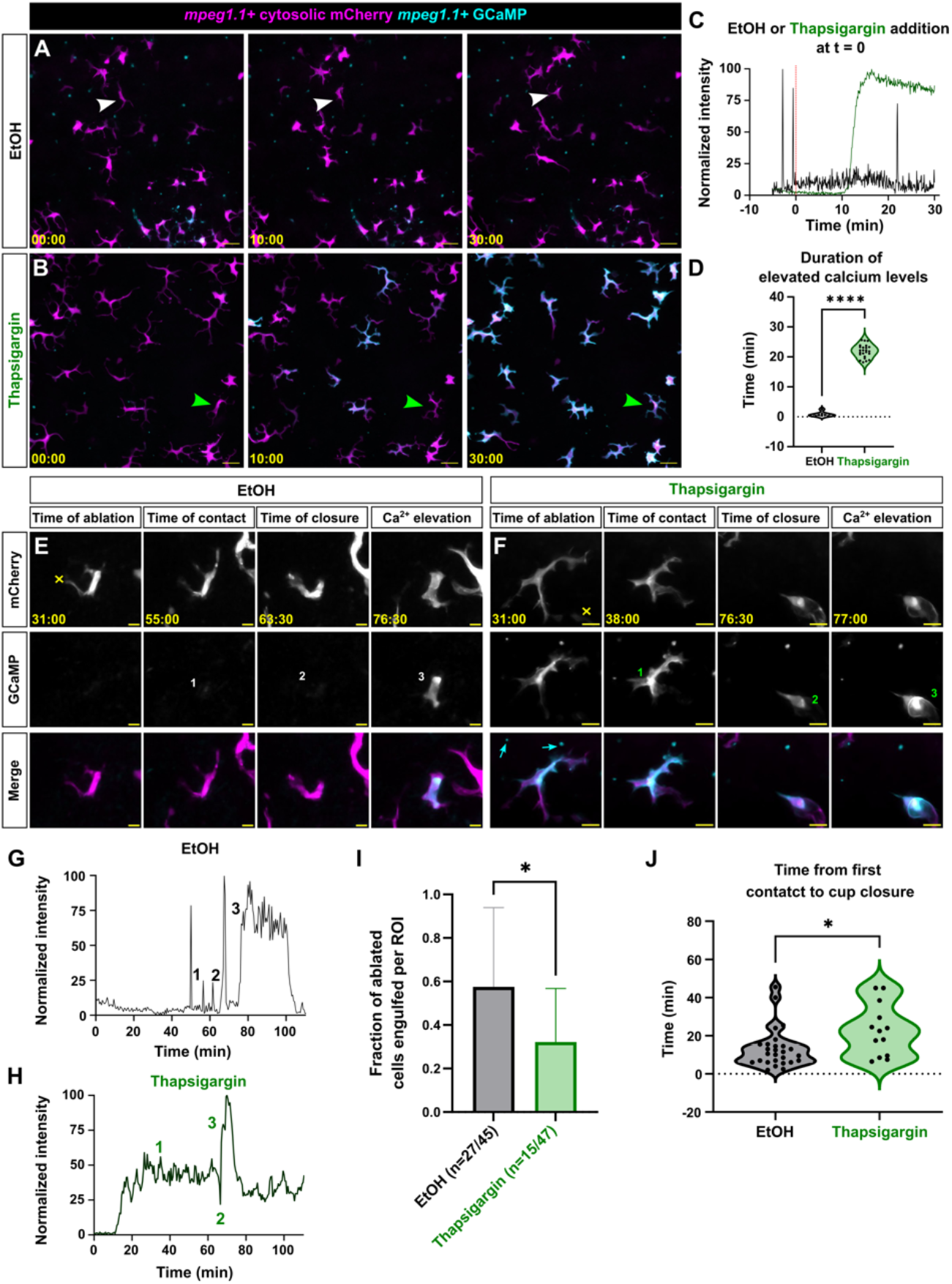
Thapsigargin-treated Langerhans cells show decreased engulfment efficiency of large debris. **A, B**. Confocal micrographs from time lapse imaging showing *Tg(mpeg1*.*1:mCherry;mpeg1*.*1:GCaMP6s-CAAX)*+ Langerhans cells during ethanol **(A)** or thapsigargin **(B)** treatment. Arrowheads indicate cells tracked in **(C). C**. Intensity plot of GCaMP signal in Langerhans cells from panels **(A, B). D**. Violin plot showing the duration of elevated calcium levels in ethanol and thapsigargin-treated Langerhans cells, *n* = 7 ethanol control cells from 3 ROIs, *n* = 24 thapsigargin-treated cells from 3 ROIs. **E, F**. Confocal micrographs from time lapse imaging showing *Tg(mpeg1*.*1:mCherry;mpeg1*.*1:GCaMP6s-CAAX)*+ Langerhans cell engulfing large debris during ethanol **(E)** or thapsigargin **(F)** treatment. Yellow X indicates the site of laser ablation. Numbers indicate relevant timepoints in **(G, H)**. Cyan arrows indicate autofluorescence. **G**. Intensity plot of GCaMP signal in Langerhans cells from **(E). H**. Intensity plot of GCaMP signal in Langerhans cells from panel **(F). I**. Quantification showing engulfment rates in EtOH and thapsigargin-treated cells from 4 (thapsigargin) or 5 (ethanol) individual experiments. **J**. Violin plots showing the amount of time from debris contact to phagocytic cup closure in EtOH and thapsigargin-treated cells, *n* = 27 ethanol-treated control cells from *N* = 5 experiments, *n* = 13 thapsigargin-treated cells from *N* = 4 individual experiments. Mann-Whitney U test was used to determine significance in **(D, I, J)**, * = *p* < 0.01, **** = p < 0.0001. Timestamps denote mm:ss. Scale bars: 20 μm (A, B, E, F).

Next, we tested if thapsigargin treatment perturbed the ability of Langerhans cells to engulf keratinocyte debris generated by laser ablation. Upon debris engulfment, ethanol-treated cells exhibited sustained elevation in calcium levels that eventually returned to baseline levels in 11/16 cells within the imaging window **(Figure 5E, G, Supplemental Video 4)**. Thapsigargin-treated cells exhibited elevated calcium levels prior to debris engulfment, and only 4/10 cells returned to baseline levels within the imaging window **(Figure 5F, H, Supplemental Video 4)**. Compared to vehicle controls, thapsigargin-treated cells exhibited a slight but significant decrease in their ability to engulf large keratinocyte debris **(Figure 5I)**. When we measured the time from debris contact to phagocytic cup closure, we found that thapsigargin-treated cells required significantly more time to close the phagocytic cup **(Figure 5J)**. Together, these data suggest that elevated calcium levels lower the efficiency of Langerhans cell phagocytic functions.

### Summary

Despite the multitude of cell types in skin, descriptions of cutaneous calcium dynamics have primarily focused on keratinocytes. Here, we directly address this gap in knowledge by analyzing the calcium dynamics of Langerhans cells, a population of tissue-resident macrophages essential for skin wound repair (Wasko et al., 2022). Using the amenability of the zebrafish scale epidermis to live-cell imaging, we found that Langerhans cells exhibited short bursts of calcium that often localized to the cell body during homeostasis. Upon engulfment of laser-ablated keratinocytes, Langerhans cells displayed increased and sustained calcium levels, sometimes lasting over an hour. This contrasts with the calcium activity in Langerhans cells that engulfed axonal debris, where we observed no such change in calcium levels. Finally, we showed that elevation of cytosolic calcium levels with a SERCA inhibitor impaired Langerhans cell engulfment of damaged keratinocytes. Previous studies implicated calcium channels in the antigen presentation functions of Langerhans cells (Diezel et al., 1989; Katoh et al., 1997) and, together with our work, suggest that calcium signaling regulates multiple dynamic, functional attributes of Langerhans cells.

Langerhans cells can most relevantly be compared with microglia, resident macrophages of the central nervous system. Ontogenetic studies suggest these two cell types share developmental origins (Kuil et al., 2020; Wang et al., 2012). Microglia also exhibit transient calcium activity during homeostasis (Umpierre et al., 2020). Prior to engulfment of myelin debris, larval microglia exhibit elevated calcium levels specifically in their dendrites (Hughes and Appel, 2020). While we did not quantify the correlation between dendritic calcium activity and likelihood of debris engulfment, we did find an increase in calcium activity in dendrites during axonal debris engulfment compared to homeostatic conditions. Recent work has shown that purinergic receptors regulate microglial calcium signaling during epileptic episodes, and interrupting this activation cascade perturbs neuronal soma engulfment (Umpierre et al., 2023). Future studies aimed at identifying the upstream molecular mechanisms that initiate increases in Langerhans cell calcium levels will shed light on damage signals sensed by immune cells within the skin microenvironment.

## MATERIALS AND METHODS

### Zebrafish

Zebrafish were housed at 26-27°C on a 14/10 h light cycle. Strains used in this study: AB, *Tg(mpeg1*.*1:GCaMP6s-CAAX)*^*co65Tg*^ (Hughes and Appel, 2020), *Tg(mpeg1*.*1:mCherry)*^*gl23Tg*^ (Ellett et al., 2011), and *Tg(Tru*.*P2rx3a:LEXA-VP16,4xLEXOP-mCherry)*^*la207Tg*^ (referred to as *Tg(p2rx3a:mCherry)*) (Palanca et al., 2013). Animals aged 6-18 months of either sex were used. All zebrafish experiments were approved by the Institutional Animal Care and Use Committee at the University of Washington (Protocol #4439-01).

### Scale removal

For scale removal, adult fish were anesthetized in system water containing 200 µg/ml buffered tricaine, and forceps were used to remove individual scales. Following scale removal, animals were recovered in system water.

### Microscopy and live imaging

An upright Nikon Ni-E A1R MP+ confocal microscope was used for all experiments. A 25× water dipping objective (1.1 NA) was routinely used. Unless otherwise stated, scales were removed and placed onto dry 6 mm plastic dishes, epidermis side up, and allowed to adhere for 30 seconds before adding L-15 medium pre-warmed to room temperature. For experiments involving axon degeneration, scales were incubated at 26°C for 90-120 min followed by imaging, which was performed at room temperature (23°C).

### Laser-induced cell damage

For laser-induced cell damage, scales were mounted into the imaging chamber as described above. Target cells at least 1 cell distance away from a Langerhans cell (∼5-15 μm) and within the same z-plane were located and ablated using a UGA-42 Caliburn pulsed 532 nm laser (Rapp OptoElectronic). The laser was focused through a 25× objective at 4× zoom. Ablation was produced in the focal plane using 15-20% power at a single point within a nucleus, firing 3 times for 3 seconds each using a custom NIS-Elements macro.

### Chemical treatments

For Thapsigargin treatments, scales were removed from fish and immediately placed in 5 mL of L-15 media. Scales were imaged for 10 minutes before careful addition of Thapsigargin to the dish while on the microscope stage. The final concentration of Thapsigargin used was 10 µM, with the appropriate ethanol vehicle control used at %v/v.

### Image analysis

#### GCaMP intensity

To quantify GCaMP intensity within Langerhans cells, the ImageJ TrackMate plugin (Ershov et al., 2022; Tinevez et al., 2017) was used to track individual cells within a given ROI over the 15-minute imaging window. Within TrackMate, the “Thresholding detector” was used to create objects around individual cells by auto-thresholding the *mpeg1*.*1*:mCherry signal, and the “Overlap tracker” was used to track the area and GCaMP sum intensity within each object over time. Average GCaMP intensity within each cell was calculated, and intensity was normalized to the highest value with each cell using GraphPad/Prism. A normalized intensity measuring greater than or equal to 25% was considered as “elevated”, and these values were used to quantify calcium activity duration and frequency. Images were manually inspected and corrected to ensure that cells that did not have elevated calcium were not included in the quantification.The total number of active cells and total number of cells per ROI were recorded and percentage of cells experiencing elevated calcium levels was calculated.

#### Location of GCaMP activity

To quantify activity location, ImageJ was used to observe each calcium elevation event as either localized to a dendrite only or to the entire cell for every time period where a given cell’s normalized intensity was equal to or above 25%. Whole cell activity includes any events that occur in multiple parts of a cell, while dendrite-only events were restricted to activity only occurring in a singular dendrite.

#### Neighborhood calcium activity analysis

To assess the correlation between neighboring Langerhans cells’ calcium activities, Pearson correlation values were calculated between all pairs of Langerhans cells in a given ROI using their normalized GCaMP intensities over time. Next, distances between the cell bodies of each pair of cells was measured in ImageJ. A graph of correlation values vs distances was produced.

#### Debris engulfment

To quantify time of debris contact to phagocytic cup closure, the transmitted light was referenced to identify when dendrites contacted laser-ablated debris. The time difference from contact to cup closure was calculated and recorded. To quantify engulfment rate, the ratio of the number of cells engulfed to the number of cells ablated in each ROI was calculated at the conclusion of the timelapse.

### Statistical analysis

Graphpad Prism was used to generate graphs and perform statistical tests. At least three individual biological experiments were performed unless otherwise noted. Statistical tests used for analysis and numbers of cells/ROIs analyzed are mentioned in the corresponding figure legend. Statistical significance was defined as * = *p* < 0.05, * * = *p* < 0.01, *** = *p* < 0.001, * * * * = *p* < 0.0001.

## ACKNOWLEDGEMENTS

We thank the LSB Aquatics staff for animal care, Dan Fong and Wai Pang Chan for imaging support, and the Appel Lab (University of Colorado, Anschutz Medical Campus) for sharing *Tg(mpeg1*.*1:GCaMP6s-CAAX)*. The authors are grateful to all members of the Rasmussen laboratory for discussion, technical assistance and continuous support. This work was supported by a postdoctoral fellowship from the Washington Research Foundation (to E.P.), an award from the Fred Hutch/University of Washington/Seattle Children’s Cancer Consortium, which is funded by P30 CA015704 (to J.P.R.), grant LF-OC-24-001646 from the LEO Foundation (to J.P.R.) and funds from the University of Washington (to J.P.R.). P.A.L.G. received support from the University of Washington Enhancing Neuroscience Diversity through Undergraduate Research Education Experiences (UW-ENDURE) program, which is funded by R25 NS114097.

## AUTHOR CONTRIBUTIONS

Conceptualization, P.A.L.G, E.P., J.P.R.; methodology, P.A.L.G, E.P., J.P.R.; formal analysis, P.A.L.G.; investigation, P.A.L.G, E.P.; resources, J.P.R.; writing – original draft, P.A.L.G, E.P.; writing – review & editing, P.A.L.G, E.P. and J.P.R.; visualization, P.A.L.G, E.P., J.P.R.; supervision, E.P., J.P.R.; project administration, E.P., J.P.R.; funding acquisition, P.A.L.G, E.P., J.P.R.

## SUPPLEMENTAL VIDEO LEGENDS

**Supplemental Video 1. Spontaneous and transient calcium activity in *ex vivo* Langerhans cells**. Maximum projection of time-lapse confocal imaging showing *Tg(mpeg1*.*1:mCherry;mpeg1*.*1:GCaMP6s-CAAX)*+ Langerhans cells exhibiting rapid and transient spikes in calcium activity. Scale bar: 20 μm.

**Supplemental Video 2. Calcium activity localized to Langerhans cell body or dendrite**. Maximum projection of time-lapse confocal imaging showing representative examples of *Tg(mpeg1*.*1:mCherry;mpeg1*.*1:GCaMP6s-CAAX)*+ Langerhans cells exhibiting spikes in calcium activity in cell body or in dendrite. Scale bar: 20 μm.

**Supplemental Video 3. Sustained, elevated calcium levels in Langerhans cell following engulfment of large debris**. Maximum projection of time-lapse confocal imaging showing *Tg(mpeg1*.*1:mCherry;mpeg1*.*1:GCaMP6s-CAAX)*+ Langerhans cell engulfing large debris following laser debris and exhibiting sustained elevation of calcium levels. Scale bar: 20 μm.

**Supplemental Video 4. Thapsigargin treatment of Langerhans cell slows engulfment of large debris**. Maximum projection of time-lapse confocal imaging showing ethanol and thapsigargin-treated *Tg(mpeg1*.*1:mCherry;mpeg1*.*1:GCaMP6s-CAAX)*+ Langerhans cells engulfing debris generated by laser ablation. Scale bar: 20 μm.

